# The *in situ* structure of Parkinson’s disease-linked LRRK2

**DOI:** 10.1101/837203

**Authors:** Reika Watanabe, Robert Buschauer, Jan Böhning, Martina Audagnotto, Keren Lasker, Tsan Wen Lu, Daniela Boassa, Susan Taylor, Elizabeth Villa

**Author notes:** These authors contributed equally (Reika Watanabe, Robert Buschauer, Jan Böhning).

## Abstract

Mutations in leucine-rich repeat kinase 2 (LRRK2) are the most frequent cause of familial Parkinson’s disease. LRRK2 is a multi-domain protein containing a kinase and GTPase. Using *in situ* cryo-electron tomography and subtomogram averaging, we reveal a 14-Å structure of LRRK2 bearing a pathogenic mutation that oligomerizes as a right-handed double-helix around microtubules, which are left-handed. Using integrative modeling, we determine the architecture of LRRK2, showing that the GTPase points towards the microtubule, while the kinase is exposed to the cytoplasm. We identify two oligomerization interfaces mediated by non-catalytic domains. Mutation of one of these abolishes LRRK2 microtubule-association. Our work demonstrates the power of cryo-electron tomography to obtain structures of previously unsolved proteins in their cellular environment and provides insights into LRRK2 function and pathogenicity.

---

Parkinson’s disease (PD) is the second most common neurodegenerative disease. About 10% of patients show a clear family history and mutations in the leucine-rich repeat kinase 2 (LRRK2) are the most frequent cause of familial PD (1, 2). LRRK2 is a 286 kDa protein with an amino-terminal half containing repeating motifs and a carboxy-terminal half composed of kinase and GTPase domains surrounded by protein-protein interaction domains (3). Most of the pathogenic mutations are found in these two catalytic domains, but all result in hyperactivation of the kinase (1, 2). Hyperactivation of LRRK2’s kinase is also reported in non-familial PD cases (4), suggesting that suppression of kinase activity might be beneficial for a wide range of PD patients.

A major barrier to understanding the mechanism of LRRK2 is the lack of structural information for the protein, which to date only includes structures of domains, either single or in tandem. Many of these are from a distantly related bacterial homolog (5–7), two are single domains from the human protein (8, 9), and two are electron microscopy (EM) structures, whose resolution was too low for molecular models to be built (6, 10). LRRK2 is involved in multiple cellular processes, including neurite outgrowth and synaptic morphogenesis, membrane trafficking, autophagy, and protein synthesis (3). LRRK2 phosphorylates a subgroup of Rab GT-Pases and modulates their membrane localization and function (11). LRRK2 forms filamentous structures surrounding microtubules (MTs) in cells, and most of the major mutations associated with familial PD enhance filament formation (12, 13). Pharmacological inhibition of LRRK2’s kinase also causes reversible recruitment of both wild-type and mutant LRRK2 to MTs (13, 14), emphasizing the need for detailed structural information of LRRK2 bound to MTs to design effective PD therapeutics.

To determine the structure of MT-associated LRRK2, we used cryo-electron tomography (cryo-ET) to image LRRK2 filaments *in situ* (Fig. 1). We grew HEK-293T human cells expressing LRRK2 bearing the pathogenic I2020T mutation tagged with YFP (referred to as “LRRK2” here) on EM grids. After vitrification, cells containing filamentous LRRK2 were identified by cryo-fluorescence microscopy. Cryo-focused ion beam (cryo-FIB) milling was then used to prepare thin slices for cryo-ET (Fig. 1A-C). LRRK2-decorated MTs were identified by correlative light and electron microscopy (CLEM) as regions where MT bundles in the transmission electron microscopy (TEM) images overlapped with the YFP signal of LRRK2 in the fluorescence images (Fig. 1D-F). Subsequently, cryo-ET was performed in these regions (Fig. 1G-I).

**Fig. 1.**
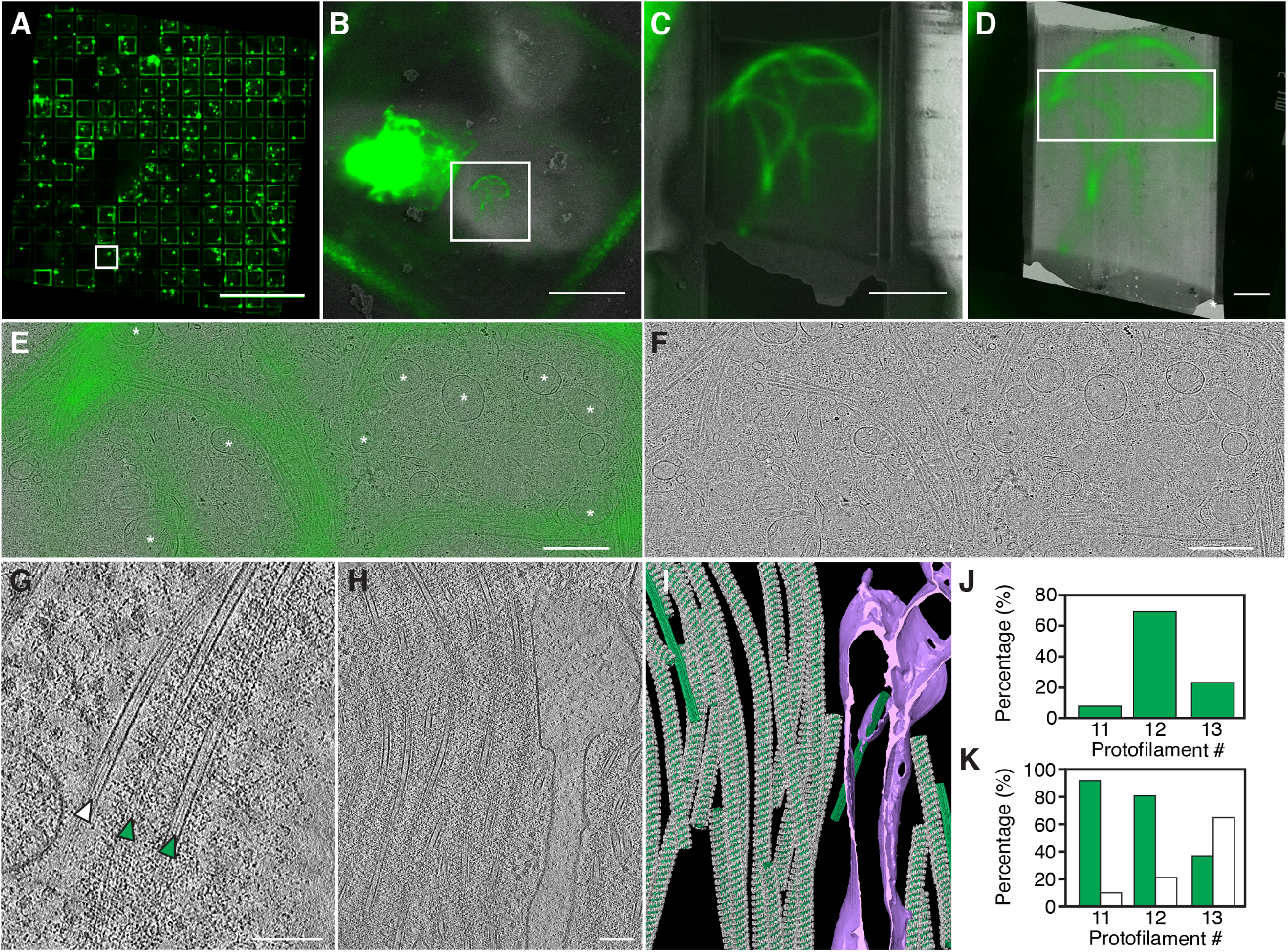
Correlative light and electron microscopy of mutant LRRK2-decorated microtubule bundles. (A) Fluorescence micrograph (FM) of a grid with HEK-293T cells expressing YFP-tagged LRRK2(I2020T) (shown in green). (B) Overlay of the fluorescence signal (green) corresponding to YFP-LRRK2(I2020T) and cryo-SEM image (greyscale) of the area marked in A. (C) Overlay of the initial FM image and an SEM image after FIB-milling of the area marked in B. (D) Overlay of the initial FM image (green) and cryo-TEM image of the area marked in B. (E and F) Enlarged TEM image of the area marked in D, with (E) and without (F) the overlaid FM image. The asterisks indicate mitochondria, which appears fragmented. (G) A slice of a tomogram showing non-decorated (white arrows) and LRRK2-decorated (green arrows) MTs. (H) Slice of a tomogram showing LRRK2-decorated MT bundle. (I) Annotated undecorated (green) and decorated (green with LRRK2 in gray) MTs and membranes (purple) in the tomogram in I. (J) Distribution of decorated microtubules with various protofilament numbers observed in LRRK2(I2020T)-expressing cells. (K) Percentage of decorated (green bar) and undecorated (white) MTs of the three different classes. Scale bars: 500 *μ*m (A), 20*μ*m (B), 5 *μ*m (C), 2*μ*m (D), 1*μ*m (E, F), 100 nm (G,H).

We first characterized the MTs in our cryo-ET tomograms. MTs are formed from heterodimers of alpha- and beta-tubulin and polymerize into polar left-handed hollow tubes, which in human cells typically contain 13 protofilaments (15). We annotated each MT individually by computationally determining its coordinates (Fig. S1A). Within bundles, most of the decorated MTs were aligned parallel to each other (Fig. 1G-I), with a center-to-center distance of 70±18 nm (Fig. S2A). This regular distribution of angles and distances suggests that LRRK2 mediates MT bundling. Most MTs were either fully decorated or not decorated, and these two populations coexisted within the same cellular regions (Fig. 1G). Protofilament number and polarity for each MT was determined by subtomogram averaging (Fig. S1B). MTs within a given bundle largely had the same polarity (Fig. S2B,C). Among LRRK2-decorated MT, we observed significant populations of non-conventional 11- (7.5%) and 12- (69.6%) protofilament MTs, in addition to the 13-protofilament MTs (22.8%) that are more typical in human cells (Fig. 1J) (15). Although we used Taxol to increase the percentage of cells showing LRRK2 filamentous structures (12), Taxol t reatment i n the absence of over-expression of LRRK2 did not alter protofilament number and showed a significantly different distance and angle distribution of MTs within bundles (Fig. S3).

Next, we used subtomogram averaging to resolve the helical arrangement of LRRK2 bound to MTs (Fig. 2A and Fig. S1C,D). We found that LRRK2 forms a right-handed double helix around MTs with all three protofilament numbers (Fig. 2A). Thus, there is a symmetry mismatch between the LRRK2 helix and the MT, which is a left-handed helix (Fig. 2A,B; helical parameters in table S1). Along the LRRK2 helix, we identified repeating units (protomers) with a C2 symmetry axis perpendicular to the filament axis. The number of LRRK2 protomers per turn matches the MT protofilament n umber, w ith 11, 12 and 13 LRRK2 protomers per turn around 11-, 12- and 13-protofilament MTs, respectively (Fig. 2A). This suggests some conformational flexibility in LRRK2 to accommodate different MT geometries. Despite this apparent flexibility, LRRK2 showed a clear preference for the smaller, and less common 11- and 12-protofilament MTs, which were almost exclusively decorated (90% and 79%, respectively; Fig. 1K). This was in contrast to 13-protofilament MTs, where only a minority was decorated (36%; Fig 1K). Furthermore, LRRK2 bound to 11-protofilament MTs appeared better resolved than LRRK2 bound to 13-protofilament MTs, despite the smaller dataset for LRRK2 bound to 11-protofilament MTs (Fig. 2A), suggesting that LRRK2 forms a more stable helix around these smaller MTs. While the physiological significance of this observation remains to be determined, unconventional MT geometries in neurons have been observed (15, 16). Since loss of dopaminergic (DA) neurons is the hallmark of PD pathology, DA neurons could have MTs predisposed to be stably decorated by the pathogenic filament-forming mutants of LRRK2. This stable decoration, even if sparse could enhance activity of LRRK2 or disturb microtubule integrity and axonal transport (17) enough to compromise cellular integrity. To determine the three-dimensional architecture of LRRK2 in the MT-associated filaments, w e p erformed further subtomogram-averaging by extracting individual protomer subtomograms from 12-protofilament MT-bound LRRK2 as this was the most abundant species in our dataset (Fig. 2A, Fig. S1E-M). We obtained a 14-Å resolution map (Fig. 2C, gold-standard FSC in Fig. S1K). Local resolution analysis (Fig. 2D) showed the region corresponding to the protomer to have resolutions of ~12 Å. To date, the only *in situ* structures of comparable resolution are of ribosomes (18, 19) and proteasomes (20), whose structures had previously been determined by other methods.

**Fig. 2.**
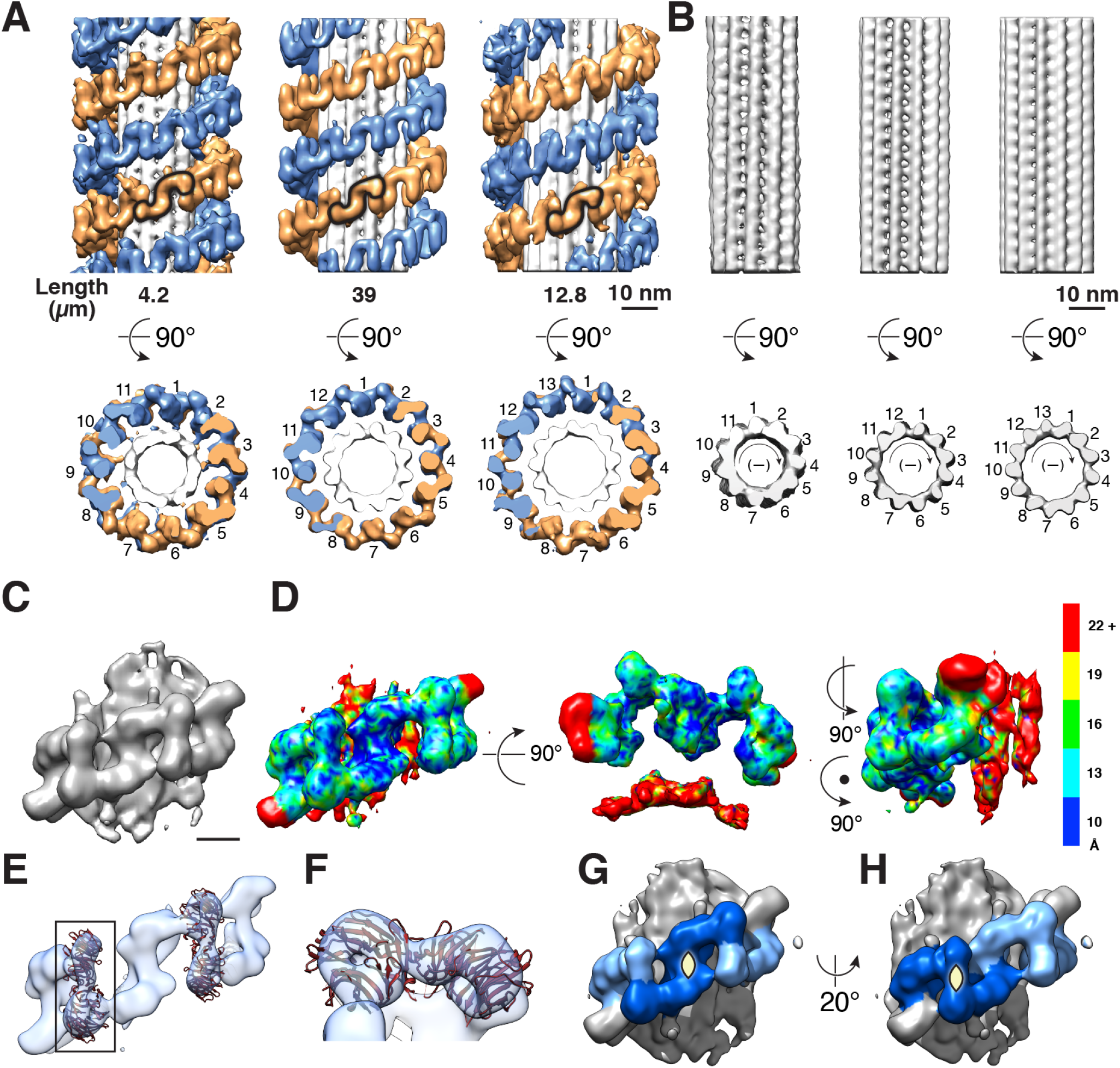
LRRK2 forms double-stranded right-handed helices around MTs with different protofilament numbers. (A) Structures of LRRK2 bound to MTs composed of 11, 12, and 13 protofilaments. The S-shaped protomer is highlighted in each reconstruction by a black outline. The total length of MTs used for averaging (in *μ*m) in shown below each MT type. The end-on views highlight that the number of protomers matches the number of protofilaments in each MT type. (B) Structure of the MTs shown in A after excluding LRRK2 with a mask. The polarity of microtubules in B is indicated with (−), meaning that the plus end of the microtubule is pointing into the page, and the minus end towards the reader. (C) LRRK2 dimer density determined by subtomogram averaging (Fig. S1E-K). (D) Local resolution of the map in C determined by ResMap. (E) The same subtomogram average shown in C is displayed at a higher threshold to highlight the fit of the known crystal structure of a LRRK2’s WD40 dimer into the density. (F) Close up of a WD40 dimer showing the hole at the center of the beta propeller. (G and H) Location of two different 2-fold symmetry axis in the LRRK2 filament (both perpendicular to the filament’s axis). The first one (G) is at the center of the protomer identified in A, while the second (H) lies at the WD40–WD40 interface.

A salient feature in our map was the C-terminal WD40 domain of LRRK2, with the hole in the beta propeller being visible at higher thresholds (Fig. 2E,F). This established that the repeating unit corresponds to a LRRK2 dimer, and can be defined as two different symmetric protomers (Fig. 2A, G). Thus, MT-associated LRRK2 filaments are formed by two homotypic interactions: a yet-to-be-determined interface within the protomer we defined (Fig. 2G) and a WD40:WD40 interface (Fig. 2H). Notably, a recent X-ray structure of a dimeric isolated human LRRK2 WD40 domain (9) fits unequivocally into our cryo-ET map of MT-bound LRRK2 (Fig. 2E, F). We could therefore use the position and orientation of the WD40 domain to determine the position of the neighboring kinase domain, which is connected to the WD40 domain through a 9 amino-acid linker. The lack of characteristic arch-shaped densities for the N-terminal Armadillo (ARM) and Ankyrin (ANK) or Leucine-rich repeat (LRR) domains in the best resolved area is consistent with the previous observation that the four C-terminal domains, namely the WD40, kinase, ROC and COR domains (Fig. 3A) are sufficient for filament formation in the cell (12).

**Fig. 3.**
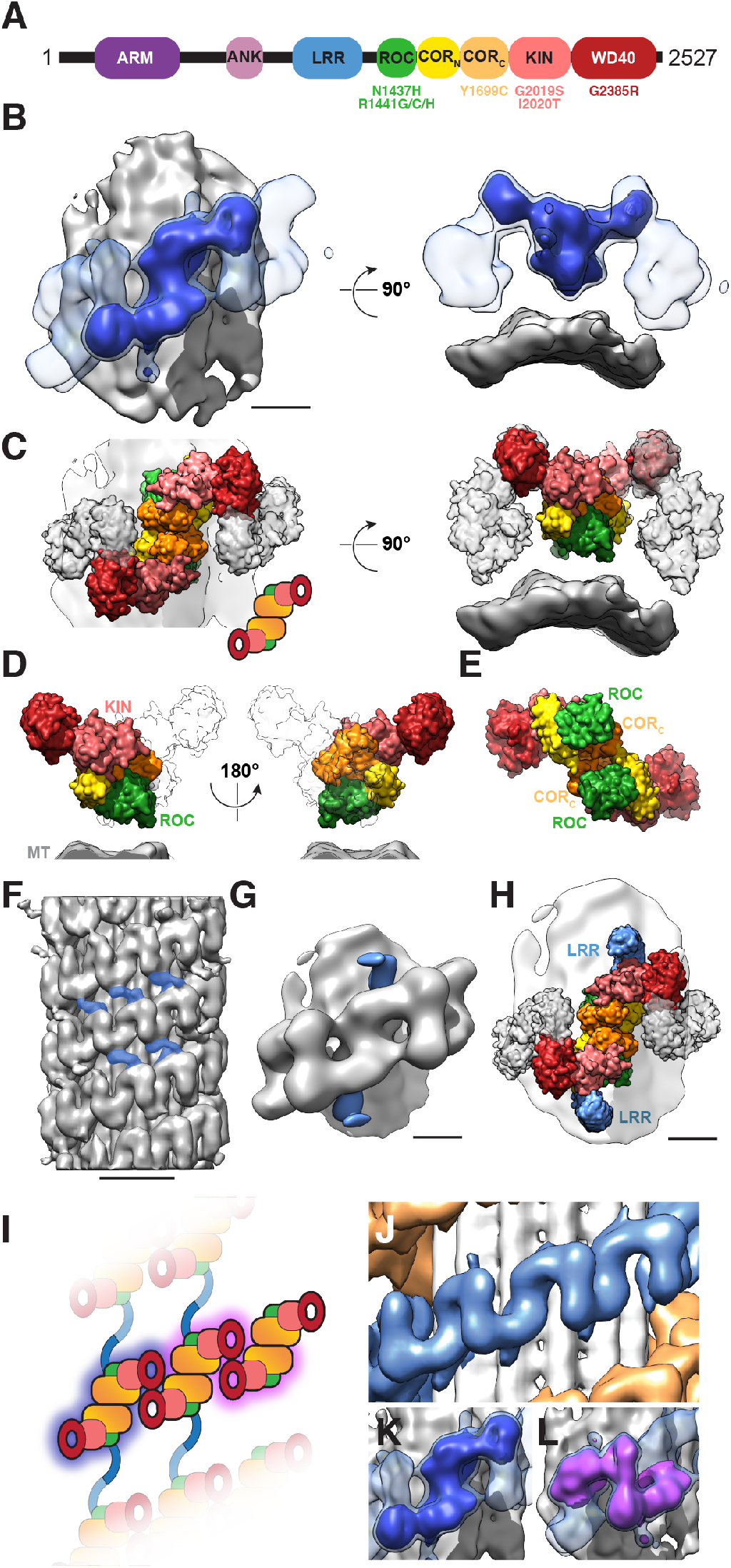
The *in situ* architecture of LRRK2 bound to MTs. (A) Schematic representation of the domain organization of LRRK2. ARM: Armadillo repeats; ANK: Ankyrin repeats; LRR: Leucine-rich repeat; ROC: Ras of complex (GTPase); CORN/CORC: C-terminal of ROC; KIN: Kinase. PD mutations are indicated. (B) The subtomogram average of the MT-bound LRRK2 filament (Fig. 2C) is shown with the protomer used for model building displayed at higher threshold (blue) within the density of the filament (semi-transparent lighter blue). (C) Molecular model of MT-bound LRRK2, shown in the same two orientations used in B. Domains are colored for the protomer (a LRRK2 dimer) with neighboring LRRK2 monomers shown in light grey. A cartoon representation of the domain architecture of the protomer is shown on the left side. (D) Two views of the molecular model for a LRRK2 monomer seen along the axis of the filament (and the MT). The second monomer in the protomer is shown as an outline. The ROC and KIN domains are labeled to highlight their location next to, and away from the MT, respectively. (E) Molecular model of the protomer viewed from the MT surface. ROC and CORC domains are labeled to highlight that dimerization is mediated by the COR_C_ domain. (F) A low-threshold isosurface of a cryo-ET map of LRRK2 bound to 12-pf MT highlights a connection between adjacent strands in the LRRK2 double helix (colored in blue). (G) Subtomogram average obtained using another alignment mask (Fig.S1I,L,M), highlighting hook-like structures (in blue) protruding from the central protomer. (H) The same molecular model for the LRRK2 protomer shown in C with a model for the LRR repeats, which were docked into the blue densities shown in G. (I-L) Architecture of the MT-bound LRRK2 filament. (I) Schematic representation of the LRRK2 filament using the same cartoon of a LRRK2 dimer introduced in C. The filament is viewed in a direction perpendicular to its axis (and that of the MT). The blue lines connecting adjacent strands represent the LRR domains. (J) Close-up of the middle map in Fig. 2A showing a stretch of LRRK2 filament corresponding to the schematic representation in I. (K, L) The two possible LRRK2 dimers – CORC-mediated in blue (K) and WD40-mediated in purple (L) – are shown in solid colors inside the semi-transparent density for the filament, and their location within the cartoon in I is highlighted by contours of the same color. Scale bars: 5 nm (B, G, H); 20 nm (F).

With the WD40 domain as an anchor point, we used an integrative modeling approach (21) (Fig. S4) to build a model of LRRK2 into the S-shaped protomer map (Fig. 3) by combining available crystallographic and homology models of the WD40, kinase, ROC and COR domains. Systematic sampling of all possible combinations of domain placements within the cryo-EM density map, subject to polypeptide connectivity and steric clashes restraints, yielded a single conFiguration of the domains of LRRK2. The structure of human LRRK2(I2020T) bound to a MT is shown in Fig. 3C-E. Our model reveals protein-protein interfaces mediated by the C-terminal of two COR domains within a protomer (Fig. 3C, E). Although our map and model are in agreement with the crystal structure of a WD40 dimer(, they differ from previous crystal structures of ROC-COR dimers from distantly-related bacterial homologs of LRRK2 (5, 22). Our model also differs from a model of an isolated inactive LRRK2 dimer in solution built on a negative-stained EM map (6). This suggests that either LRRK2 uses a different COR:COR dimerization interface from that observed in the reported structures or that conformational changes are involved in the formation of filaments.

In our structure, the ROC (GTPase) domain is closest to the MT surface (Fig. 3C,D), consistent with observations that the ROC domain of LRRK2 is sufficient for interaction with tubulin heterodimers (23). GTP binding and hydrolysis by the ROC domain have been proposed to be involved in dimerization (22), filament formation around MTs (13), and regulation of kinase activity (24). The kinase domain is distal to the MT and exposed to the cytoplasm (Fig. 3C). Since inhibitors that capture the kinase in an ATP-bound active-like state (25, 26) are known to induce filament formation (13, 14) and all other filament forming LRRK2 mutants (N1437H, R1441G/C and Y1699C) also results in hyper-kinase activity in cells (11–13), we expect that our structure captures an active kinase conformation, which may be facilitated through oligomerization. Taken together, our data suggests that MT-binding plays a scaffolding role in the oligomerization and activation of LRRK2.

Outside the protomer, our tomograms showed densities connecting the two strands of the LRRK2 helices that decorate MTs, suggesting inter-helix contacts (Fig. 3F). We generated a 18-Å resolution map encompassing a larger volume around the protomer that revealed two hook-like structures stemming from the protomer (Fig. 3G, Fig. S1I,L,M, Fig. S5). Given their shape and adjacency to the ROC domain, we propose that these densities likely represent the LRR domains, (Fig. 3H). Thus, there is a third homotypic interaction, mediated by the LRR domains, that connects the strands of the double helix, which may determine its regular spacing along the MT (Fig. 3F). In our density maps, the N-terminal ARM and ANK domains were not resolved, presumably due to their flexibility. H owever, w e o bserved u ndefined de nsity in the periphery of the LRRK2-decorated MTs but not in undecorated MTs (Fig. S6A, B). The density approximately spans the distance between decorated MTs within a bundle, suggesting that the ARM and ANK domains may form connections between adjacent MTs.

Finally, to test our model we set out to determine if disrupting one of the two dimer interfaces we observed affected the formation of MT-associated LRRK2 filaments i n cells. We focused on the LRRK2 WD40:WD40 interface because a mutation, G2385R, in the WD40 domain is a risk factor for PD (27, 28) and has been shown to disrupt WD40 dimer formation *in vitro* (9). G2385 is located at the WD40:WD40 interface between protomers (Fig 4A, B). We hypothesized that disrupting this interface would lead to defects in filament formation. To test this, we expressed Halo-tagged wild-type or G2385R LRRK2 in human HEK-293T cells in the presence or absence of the LRRK2-specific kinase inhibitor MLi-2 (29), which has been shown to induce LRRK2 filament formation (7, 13). We found that MLi-2 induced filament formation in cells expressing wild-type LRRK2, but not in cells expressing G2385R LRRK2 (Fig. 4C, D) as shown previously using another LRRK2 kinase inhibitor, LRRK2-IN-1 (30). These data support our model that the WD40:WD40 protomer interface is important for the formation of LRRK2 filaments. Given the correlation between breaking the dimerization interface, loss of filament formation, and change in kinase activity seen in the G2385 LRRK2 mutant (9, 11, 30–32), supporting that oligomerization plays a role in regulating LRRK2 kinase activity.

**Fig. 4.**
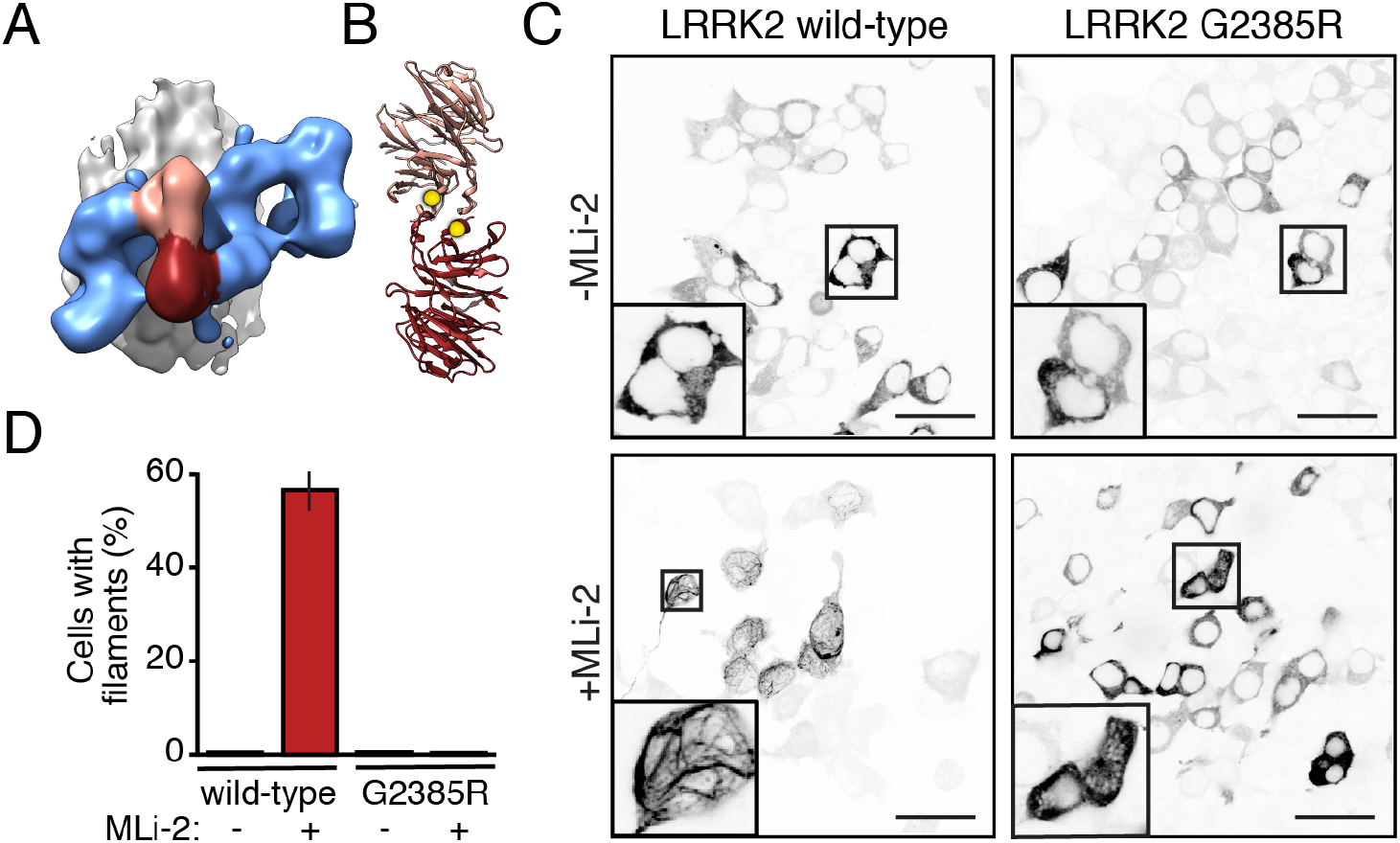
The WD40:WD40 interface is necessary for LRRK2 filament formation. (A) The subtomogram average of the MT-bound LRRK2 filaments is shown with the MT in grey, the LRRK2 filament in blue, and the WD40-WD40 dimer connecting two protomers in pink/maroon. (B) The crystal structure of the WD40 dimer with monomers colored as in A, and the G2385 residues at the interface shown as yellow spheres. (C) HEK-293T cells transiently expressing Halo-tagged LRRK2 were labeled with TMR-ligand. Representative images are shown for cells expressing wild-type (left panels) and G2385R (right panels) LRRK2, incubated in the absence (top) and presence (bottom) of 100 nM MLi-2, a LRRK2 kinase inhibitor, for 2 hours. The contrast was inverted for better visualization of the LRRK2 signal where black is the highest signal and white is no signal, i.e. background. (D) Percentage of transfected cells showing LRRK2 filament formation, defined by bright rod-like filamentous structures. Data represent three independent experiments. The total number of transfected cells considered in the analysis was 1705 for wild-type/-MLi-2; 1374 for wild-type/+MLi-2; 1464 for G2385R/-MLi-2; 1472 for G2385R/+MLi-2.

Here, we have shown that recent advances in cryo-ET can be used to determine novel structures of proteins *in situ*, making it possible to generate structural hypotheses in a cellular context. We determined the first structure of full-length human LRRK2 and its association with MTs. We also show the power of integrative modeling to extract structures from *in situ* cryo-ET data. Since well-characterized LRRK2 kinase inhibitors enhance the ability of LRRK2 to form filaments around MTs (13, 14), the structure presented here not only provides insight to understanding the pathogenic state of familial PD, but is also the first step towards designing improved LRRK2 kinase inhibitors.

## Materials and Methods

### Cell culture

HEK-293T cells were cultured in Dulbecco’s Modified Eagle’s Medium (DMEM) containing GlutaMAX-I [Thermo Fisher Scientific, (TFS)], 10% HyClone bovine calf serum (GE Healthcare) and 100 U/mL HyClone penicillin-streptomycin (GE Healthcare) at 37°C and 5% CO2.

### Plasmids

Plasmid encoding full-length LRRK2(I2020T) fused with YFP at the N-terminus was described previously (12). The construct for the N-terminus Halo-tagged full-length LRRK2 was cloned using the Gibson Assembly method. The LRRK2 gene was cloned by using the primer sets: 5’-GCGATAACATGGCTAGTGGCAGC-3’ and 5’-GGGGTTATGCTAGTTACTCAACAGATGTTCGTCTC-3’ with pENTR221-LRRK2 (Addgene #39529) as the template. The N-terminal Halo-tagged vector segment was cloned by using the primer sets: 5’-GAGTAACTAGCATAACCCCTTGGC-3’ and 5’-CACTAGCCATGTTATCGCTCTGAAAGTACAGATC-3’ with the pHTN HaloTag® CMV-neo Vector (Promega) as template. The two segments were assembled using the NEBuilder® HiFi DNA Assembly Kit following their suggested protocol. The G2358R mutation was made by using the NEB Q5-site directed mutagenesis kit, and the primers were designed based on the online tool NEBBaseChanger. The constructs were sequenced and expressed in HEK-293T cells to confirm the expression of full-length LRRK2.

### TEM grid preparation

For expression of LRRK2, the HEK-293T cells were transfected with a plasmid encoding full-length LRRK2(I2020T) fused with YFP at the N-terminus using a Lipofectamine 3000 transfection kit (Invitrogen) according to the manufacturer’s protocol. After 2 days, 5*μ*M paclitaxel (aka Taxol, form Science Signaling) was added. After 16 to 24 hours of Taxol treatment, the cells were detached, and counted with a hemocytometer. Quantifoil 200 mesh holey carbon R4/1 or R2/1 copper grids (Quantifoil Micro Tools) were glow discharged for 60 s at 0.2 mbar with 20 mA using a PELCO easiGlow glow discharge system (Ted Pella). Just before deposition of cells on TEM grids, 1.5 *μ*l of poly-lysine (Sigma) was applied and ~2000-6000 cells were deposited onto a grid by pipetting 3-10 *μ*L of detached cells onto the grid. Blotting and plunging was performed using a custom-made plunger (Max-Planck-Institute for Biochemistry) using 50/50 mixture of liquid ethane and propane (Airgas) cooled to liquid nitrogen temperature. The grids were clipped onto Autogrids (TFS) and samples were kept at liquid nitrogen temperature throughout the experiments.

### Cryo-fluorescence microscopy

For cryo-fluorescence microscopy, grids were observed with a CorrSight microscope (TFS) using EC Plan-Neofluar 5x/0.16NA, EC Plan-Neofluar 20x/0.5NA, and EC Plan-Neofluar 40x/0.9NA air objectives (Carl Zeiss Microscopy), an Oligochrome lightsource, which emits in 4 different channels (405/488/561/640 nm) (TFS) and a 1344×1024 px ORCA-Flash 4.0 camera (Hamamatsu). Acquisition and processing of the data was performed with the MAPS 2.1 software (TFS). After acquisition of a grid map at 5X magnification, regions of interest were imaged at 20x or 40x magnification, to identify cells with regions containing LRRK2 filamentous structures.

### Cryo-focused ion beam milling

Micromachining of frozen hydrated cells was performed in a Scios DualBeam FIB/SEM microscope equipped with a cryo stage (TFS). The sample chamber of the microscope was kept at a pressure below ~1×10^−6^ mbar. A low-magnification SEM image encompassing the entire grid using an acceleration voltage of 5 kV, a beam current of 25 pA, and a dwell time of 200 ns was correlated with the grid map from the cryo-light microscope using MAPS 2.1 (TFS) to identify regions of interest on TEM grids. A thick platinum layer (typically ~2*μ*m) was deposited on the sample to improve its conductivity and to reduce streaking on the lamella caused by local variations in mass density that result in variations in lamella thickness, known as curtaining (33). An integrated gas injection system (GIS) is used to deposit the precursor compound trimethyl(methylcyclopentadienyl)platinum(IV) (34, 35). To this end, the grid was placed 11.5 mm away from the GIS with a nominal stage tilt of 7°, and the GIS was opened for 13 seconds. Cells were targeted for FIB-milling that satisfied the following conditions: (1) showed a filamentous LRRK2 phenotype; (2) were located within the center of a grid square, and (3) were located on a grid square that was at most five squares from the center of the grid (the central ~1mm^2^ area of the grid). The stage was positioned at a nominal tilt of 11-18°, corresponding to a milling angle of 4-11°. FIB milling was performed in three steps with decreasing ion beam currents and a fixed acceleration voltage of 30 kV. By decreasing the current in three steps, the beam spread was reduced to a minimum at the final stages of milling, while accelerating the milling process at initial stages. For the rough milling step an ion beam current of 0.3 nA was used. The current was reduced to 0.1 nA for the intermediate step and to 30 pA for fine milling. The target lamella thickness was ~100 nm. During the milling process, lamella thickness was estimated utilizing thickness-dependent charging effects observed in the SEM images at different acceleration voltages (35). At the end of the session, the grids were transferred out under vacuum and stored in liquid nitrogen.

### Cryo-electron tomography

Tomographic tilt series were recorded in a Tecnai G2 Polara (TFS) equipped with a field emission gun operated at 300 kV, a GIF Quantum LS post-column energy filter (Gatan) and a K2 Summit 4k×4k pixel direct electron detector (Gatan). The FIB-milled grids were loaded into the TEM using modified Polara cartridges, which securely accommodate Autogrids (36). The milling slot of each grid was aligned perpendicular to the tilt axis of the microscope. This way the inherent tilt axis of the slanted lamella was parallel with the tilt axis of the microscope stage allowing separation of focus and record areas according to low dose practice along the tilt axis at equal heights. Tilt series were acquired at a target defocus of 5*μ*m and a pixel size of 2.2 or 3.5Å using the SerialEM software (37) in low-dose mode. The dose-symmetric tomography acquisition scheme (38), was modified to account for the pre-tilt of the lamella, and to allow for the acquisition to stop when the image does not yield enough counts with a target dose due to thickness, which occurs on the side of the pre-tilt first. The K2 detector was operated in counting mode and the images were divided into frames of 0.075 to 0.1 s. The tilt range was typically between ±50°and ±70°with increments of 2 and 3°and the total electron dose was around 180 e/Å2 and ∼120 e/Å^2^, respectively.

### Tomogram reconstruction and annotation

The tilt series were and aligned and dose-weighted according to the cumulative dose using MotionCor2 (39). Alignment of the tilt series and tomographic reconstructions were performed using Etomo, part of the IMOD package (40). Since no fiducial markers were present on the lamellas, tilt-series alignment was performed using patch tracking. Contrast transfer function (CTF) correction was performed in IMOD (40) using defocus values measured by CTFFIND4 (41) prior to dose-weighting via Fourier filtering. Tomograms were reconstructed using weighted back-projection. Tomograms were 4x-binned (without CTF correction) and used for segmentation of microtubules using the filament tracing function in Amira (TFS). After filament tracing, the resulting center coordinates were re-sampled equidistantly using a script written in MATLAB. For each point along the filaments, initial Euler angles were assigned for subtomogram averaging. Within a filament, angles were assigned such as the direction along the filament axis was the same for all particles. The in-plane Euler angle (orthogonal to the axis of the filament) was randomized to minimize the effect of the missing wedge.

### Subtomogram averaging

To determine the protofilament number of single microtubules, 4x binned tomograms were used for extraction of subtomograms with a box size of 38 nm along the microtubule at 4 nm spacing using the Dynamo software, and an initial average for each microtubule was created using the pre-assigned Euler angles described above. Each filament was separately aligned and averaged using Dynamo (42), using the initial average as a reference. Three iterations of translational and orientational alignment of the first two Euler angles followed by five iterations of translational alignment and orientational alignment of the third Euler angle, with the latter restricted to 18-36 degrees. Particles were low-pass filtered to 22-30 Å for alignment. 2D projections along the MT axis were calculated in MATLAB, and the 2D projections as well as the 3D averages were visually inspected in MATLAB and IMOD to determine protofilament number and plus/minus end polarity of each microtubule. Assignment of MT polarity can be achieved by inspecting a cross section of the averages from the plus end (anti-clockwise slew; Fig. S1B, left column) or from the minus end (clockwise slew; Fig. S1B, right column) respectively (43). Microtubules with unclear or ambiguous protofilament number or polarity were discarded. Then, new particles with a box length of ~70 nm were extracted from 4x binned tomograms, and for each microtubule class (11-, 12- or 13-protofilament MTs) all particles representing the respective class were submitted to alignment and averaging in Dynamo, using an initial average as a reference. In order to separate decorated MTs from non-decorated MTs, multi-reference alignment in Dynamo was conducted using two templates: (1) a LRRK2-decorated MT template (being the average from the previous step) and (2) a non-decorated MT template (being the average from the previous step with the density of LRRK2 masked out) using a hollow cylindrical classification mask corresponding to the LRRK2 density. The same alignment parameters as described above were used. In order to further improve the structure of LRRK2 bound to MTs, particles sorted to class 1 (decorated MTs) were further aligned by using a hollow cylindrical alignment mask that included LRRK2 but excluded the MT. For obtaining MT averages (Fig. 2B), all particles (decorated and undecorated) with the same protofilament c ount a nd p olarity w ere a ligned by using a cylindrical alignment mask containing just the MT region. The resulting averages were used to estimate the helical parameters of the LRRK2 double-stranded helix and MTs as detailed below. Coordinates of particles along the helical path were used to extract particles from unbinned, dose-weighted and non-CTF-corrected tomograms, and used to create an initial average using pre-assigned orientations. 3D refinement of extracted particles was performed in RELION (44, 45) to create a template for subsequent classification. This density was used as a reference for 3D classification into three classes, employing a soft mask covering the LRRK2 protomer and a regularization parameter of 2-4. Class averages were inspected in UCSF Chimera (46), and classes with high levels of noise were excluded. A density corresponding to the ring-shaped WD40 domain was clearly resolved in one class (class 1 in Fig. S1G). Particles belonging to this class were subjected to gold-standard refinement u sing t wo different masks (masks A and B; Fig. S1H,I) and post-processing without B-factor sharpening in RELION. Local-resolution maps were calculated using ResMap (44, 45). Maps and structures were visualized using UCSF Chimera (46) and VMD (47).

### Measuring distances/angles between MTs

To calculate the angles and distances between filaments (microtubules), the distances between all points sampled every 4 nm along each filament and all points on all other filaments within the tomogram were measured. The resulting distance matrix was then reduced to point pairs with distances of 100 nm or less. For each point along a filament, we calculated (1) a tangent vector along the filament at that point, considering the nearest points along the same filament, (2) the distance to the nearest point in each neighboring filament, and (3) the angle between the vectors at these two points, where an angle of zero corresponds to parallel filaments. A 2-D histogram of the distance between filaments vs angle between filaments was calculated in MATLAB.

### Determining helical parameters

Helical parameters of non-symmetrized MTs and LRRK2 helices were obtained using an autocorrelation function as implemented in Dynamo. To obtain the helical parameters for MT-bound LRRK2, averages of each were obtained by masking out the volume corresponding to the other, i.e., MT helical parameters were obtained from an average obtained with LRRK2 excluded through a mask, and vice versa.

### Integrative Modeling of LRRK2 bound to MTs

LRRK2 is a 286kDa protein formed of seven domains: armadillo (ARM), Ankyrin (ANK) and Leucin-rich repeat (LRR) domains, a Ras of complex proteins (ROC), and C-terminal of ROC domain (COR), kinase (KIN) and a WD40 domains (Fig. 3A). To determine the architecture of LRRK2/I2020T bound to MTs, we applied an integrative approach implemented in the open source Integrative Modeling Platform (IMP) package (48). The integrative modeling process consists of four stages: (1) gathering data, (2) choosing how to represent the system and translating the information into spatial restraints, (3) determining an ensemble of structures that satisfy these restraints and, (4) validating the model. This approach has been applied to determine the structure of numerous biological complexes including the 26S proteasome (49), the mediator complex (50) as well as the nuclear pore complex (51). To our knowledge, this work represents the first application of integrative modeling to solve a novel structure determined *in situ* using cryo-ET. Given the resolution of the cryo-ET map, the modeling was limited to the location and orientation of the domains within the density, treated as rigid bodies. The clear identification into the cryo-ET density map of the characteristic donut-shape of the WD40 provided the starting point for the LRRK2 domains allocation and allowed us to assess that the assignment of the LRRK2 is possible up to the last four domains (WD40, KIN, COR, ROC) while the density related to the LRR, ANK and ARM is visible only at higher threshold, presumably due to their intrinsically flexibility.

#### Stage 1: Gathering Information

Three different types of data were used for structure determination: A. Cryo-ET map of LRRK2(I2020T) protomer: a 14-Å *in situ* cryo-ET map was determined as detailed above. The protomer corresponds to a dimer LRRK2 bound to the MTs (Fig 3B). B. Atomic models of the domains composing LRRK2: X-ray crystallography structure of the human LRRK2 WD40 (9) (PDB:6DLO) Homology model of the kinase domain of human LRRK2 (Uniprot ID Q5S007, residue 1883-2135) in its active-like conformation was generated by homology modeling with SWISS-MODEL (52, 53). The crystal structure of the Roco4 Kinase domain bound to AppCp from *D. discoideum* was used as a template (54) (PDB:4F0F, homology 43%). Alignment was generated by ClustalOmega (55) (Fig. S7 and S8). A model of the monomeric ROCCOR domain of human LRRK2 protein (Uniprot ID Q5S007, residue 1332-1838 was obtained by homology modeling with SWISS-MODEL (52, 53). The structure of *C.tepidum* Roco protein (5) (PDB:3DPU) was used as template and aligned to the human LRRK2 sequence of domains ROC and COR by using ClustalOmega (55) (Fig. S7A). Normal model analysis (NMA) with Bio3D package in R (56) was performed on the ROCCOR template revealing a hinge in the COR domain (L1669-I1689) which corresponds to a missing loop area in the template PDB structure. Therefore, we split the ROCCOR domain into three separate rigid bodies namely, CORC, CORN, and ROC. (PDB:3DPU, homology 37%, 37% and 38% respectively; Fig. S7 and S8). C. Since all the domains are part of a single polypeptide chain of LRRK2, the amino acid stretches between the domains were considered as linkers connecting the N- and C-termini of consecutive domains. A worm-like-chain model (57) was used to model the connecting linkers, with an average end-to-end distance of 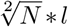, where *N* is the number of amino acids and *l* = 3.1Å is the length of one amino acid measured as the typical average distance between alpha carbons of adjacent amino acids (Fig. S8).

#### Stage 2: System Representation and Translation data into Spatial Restraints

Each domain was represented as a rigid body at atomic resolution. Five types of spatial restraints were used during monomer conFiguration and dimer refinement in Stage 3. Cryo-ET density restraint: (a) an IMP scoring term that considers the percentage of atoms that are included in the map, used both in monomer conFiguration and refinement. (b) Cross correlation o f m odel t o m ap (implemented in MDFF), used to select models in monomer conFiguration. Location of WD40: based on the characteristic donut shape in the density, the WD40 could be unequivocally located in the cryo-ET density map and thus its location was constrained to that region in the map (Fig. S8), used in monomer conFiguration. Chain connectivity restraints: distance restraints between the C-term and N-term of consecutive domains as described above (Fig. S8), used both in monomer conFiguration and dimer refinement. Excluded volume restraints: Overlap between domains is assessed as an excluded volume restraint (steric clashes of less than 10% and 5% of alpha carbons), used both in monomer conFiguration and dimer refinement r espectively. Dimer symmetry restraints: (a) Overlap between monomers was assessed using excluded volume (steric clashes of less that 10% of alpha carbon atoms between LRRK2 monomers), used in the final stage of monomer conFiguration. (b) Symmetry restraints implemented in IMP used during dimer refinement.

#### Stage 3: Ensemble Sampling

To create an ensemble of LRRK dimers that incorporates the data available and satisfies the restraints, we applied a sampling protocol consisting of two steps: monomer conFiguration and dimer refinement. A. Monomer conFiguration. To determine the monomer conFiguration we used the MultiFit module of IMP (58). First, rigid-body fitting of each of the individual five domains to a dimer protomer map were sampled at 5°resolution using colores (59). The best 10,000 fits for WD40, ROC, C ORN and CORC and 5,000 fits f or KIN were c onsidered. WD40 fits were filtered to be placed within the donut-shaped density to satisfy constraint ii. From these fits, we selected a single position and orientation that matched that of the dimer in the dimer crystal structure (9). Second, we looked at all 5000 fits of the kinase and selected those that satisfied the distance linker of 9 residues (restraint iii). Only six fits satisfied this restraint, with an RMSD of 4.7Å between them, and thus we used a single kinase fit for the subsequent s teps. N ext, the fits for ROC, C ORN and C ORC were filtered to include only those fitted to the same monomer within the protomer map, and not overlapping with the placed WD40 and kinase, yielding 1188, 1446 and 1148 models respectively. MultiFit was then used to sample all possible combinations of the ROC, CORN and CORC domain fits and scored for spatial restraints i(a), iii and iv. A total of three million models were initially scored. From that, we filtered the solutions based on respecting the N-C and having not major compenetration between domains, (corresponding to an EV score of <100,000). This yielded 786,211 models. Third, the ensemble was clustered based on RMSD of 8 Å, leading to 256 clusters, that could be further classified into models with different architectures, e.g., swaps between ROC, CORN and CORC domains. Next, the models were filtered to allow only clashes of up to 10% of the alpha carbons within domains as measured in VMD (47). From here, only two clusters remained, one with 31,140 models, and the other with 394 models. Fourth, dimers were built from each monomer by applying a symmetry transformation. All models in the second cluster had major clashes at the dimer interface. Thus, only one ensemble of 31,140 models remained corresponding to the same domain architecture. The ensemble was filtered to satisfy restraints i, iii, iv, and v(a), resulting in 1,137 models. B. Dimer refinement. We generated an ensemble of LRRK2 dimers by rigid-body fitting each monomer model into the dimer density map, as detailed above. We then clustered the dimers ensemble into 48 clusters based on an RMSD of 8 Å. From each cluster, we refined a representative dimer using a simulated-annealing Monte-Carlo (MC) protocol implemented in IMP. During the refinement, local perturbations in positions and orientations of the ten rigid bodies were sampled and assessed using a scoring function including terms i-v. Minima from the refinement protocol were gathered and used in the refined ensemble assessment (stage 4).

#### Stage 4: Assessment of Data and Structure

Our extensive sampling rendered a single architecture of LRR2 dimers. In order to validate this structure, we first looked at the overall conformation and its congruency with other available data. We found that our model agrees with the following independent observations: The LRR domain fits into the density and is in the proximity of the ROC domain (Fig. 3H). The LRR model corresponds to residues 129-444 of the LRR in a crystal structure of a homologous Roco protein (PDB ID: 6HLU), and was fitted using the rigid-body fitting function of Chimera (46). The ROC domain is proximal to the microtubules, consistent with observations that the ROC domain of LRRK2 is sufficient for interaction with tubulin heterodimers (23). The dimerization of the WD40 observed in a recent crystal structure (9) matches the architecture in our model, and is consistent with our data showing no filament formation (Fig. 4). Although we used the crystal structure to assign the WD40 conformation, a preliminary global alignment using MultiFit (data not shown) confirmed this to be the best location of the WD40 independently of the crystal structure. The monomer ensemble obtained was analyzed to assess the uncertainty on the positions and orientations of each domain (Fig. S9). The analysis on the domains variability assessed a narrow conformational space visited by COR, CORC and a broader conformational space visited by ROC (Fig. S9). During refinement, each model improved its conformation in terms of the restraints, without major rearrangements. Thus, the ensemble reflects the variability of the structure given the available experimental data. Altogether, we determined the architecture of LRRK2, with certainty in the location of the domains, and varying uncertainty in their orientation, as expected from the resolution of our data.

### Confocal fluorescence microscopy and statistical analysis

For transient protein expression of N-terminally Halo-tagged full-length wild-type and G2385R LRRK2, HEK-293T cells were plated onto 6-well dishes containing poly-D-lysine-coated glass coverslips or onto poly-D-lysine-coated glass-bottom culture dishes (MatTek Corporation, Ashland, MA, USA, Cat. No. P35GC-0Ð14-C) and allowed to grow to 40-50% confluency. Cells were transfected using the Lipofectamine 2000 reagent (ThermoFisher Scientific, USA) according to manufacturer’s protocol using 1 μg of tagged Halo-LRRK2 DNAs. After 48 hours, cells were treated with the LRRK2 inhibitor MLi-2 (100 nM) for 2 hours and then labelled with the TMR ligand (3 *μ*M)(Promega). After 15 minutes, cells were washed with fresh media and then fixed with pre-warmed (37°C) 4% paraformaldehyde in phosphate-buffered saline (PBS) for 15 minutes at room temperature. Cells were then washed in PBS and mounted with the in-house made antifade agent gelvatol. Confocal imaging was performed with the inverted Olympus Fluoview 1000 laser scanning confocal microscope using a 561 nm laser line and a 60X oil immersion objective lens with numerical aperture 1.42. Z-stack images were acquired with a step size of 0.3 microns and processed using the Fiji software package (60). Cells expressing the different LRRK2 mutants were assessed for the presence of clear filamentous structures and quantified in three independent experiments.

## ACKNOWLEDGEMENTS

We thank Daniel Castano Diez and Paula Perez Navarro for assistance with Dynamo; Guillaume Castillon, Alexis Rohou, Junru Hu, and members of the Villa lab for stimulating discussions and technical support. This work was supported by an NIH Director’s New Innovator Award 1DP2GM123494-01 (to E.V.), a Michael J. Fox Foundation grant (to S.T., E.V and D.B.), and a Branfman Family Foundation award (to D.B). M.A. is supported by a postdoctoral fellowship from the Visible Molecular Cell Consortium at UC San Diego. T.L. is supported by a Taiwan MOE fellowship. We acknowledge the use of the UC San Diego cryo-Electron Microscopy Facility (partially supported by a gift from the Agouron Institute to UC San Diego), the NIH National Center for Microscopy and Imaging Research (P41GM103412), and the San Diego Nanotechnology Infrastructure (SDNI) of UC San Diego, a member of the National Nanotechnology Coordinated Infrastructure, which is supported by the National Science Foundation (ECCS-1542148). We thank the Henriques Lab for the bioRxiv LATEX template.

## Supplementary Data

### Effects of Taxol

To examine the effect of Taxol incubation (5 *μ*M) performed to increase the percentage of cells showing LRRK2 filamentous structure (12), MTs in the HEK-293T cells incubated with Taxol, in the absence of LRRK2 overexpression, were similarly analyzed (Fig. S3). In Taxol-treated HEK-293T cells, MTs were less clustered compared to LRRK2 decorated MTs (Fig. S3A, B). The center-to-center distance between MT was the average distance of 47±32 nm (peak 31 nm) (Fig. S3C). The wider distribution compared to LRRK2 expressing Taxol-treated cells (Fig. S2) indicates more random MT arrangements in the Taxol-treated cell suggesting that LRRK2 itself mediates MT bundle formation observed in Fig.1H, I and Fig. S2. The 31 nm peak distance likely corresponds to the closest possible packing (non-decorated 13-protofilament MT diameter is 25 nm; Fig. S3A-C). In these cells, 8% of the total MTs belonged to 12-protofilament MTs, and no 11-protofilament MTs were found, while 92% of MTs observed were 13-protofilament MTs suggesting that Taxol treatment used in our experiments did not drastically affect MTs population in HEK-293T cells (Fig. S3D).

---

**Fig. S1.**
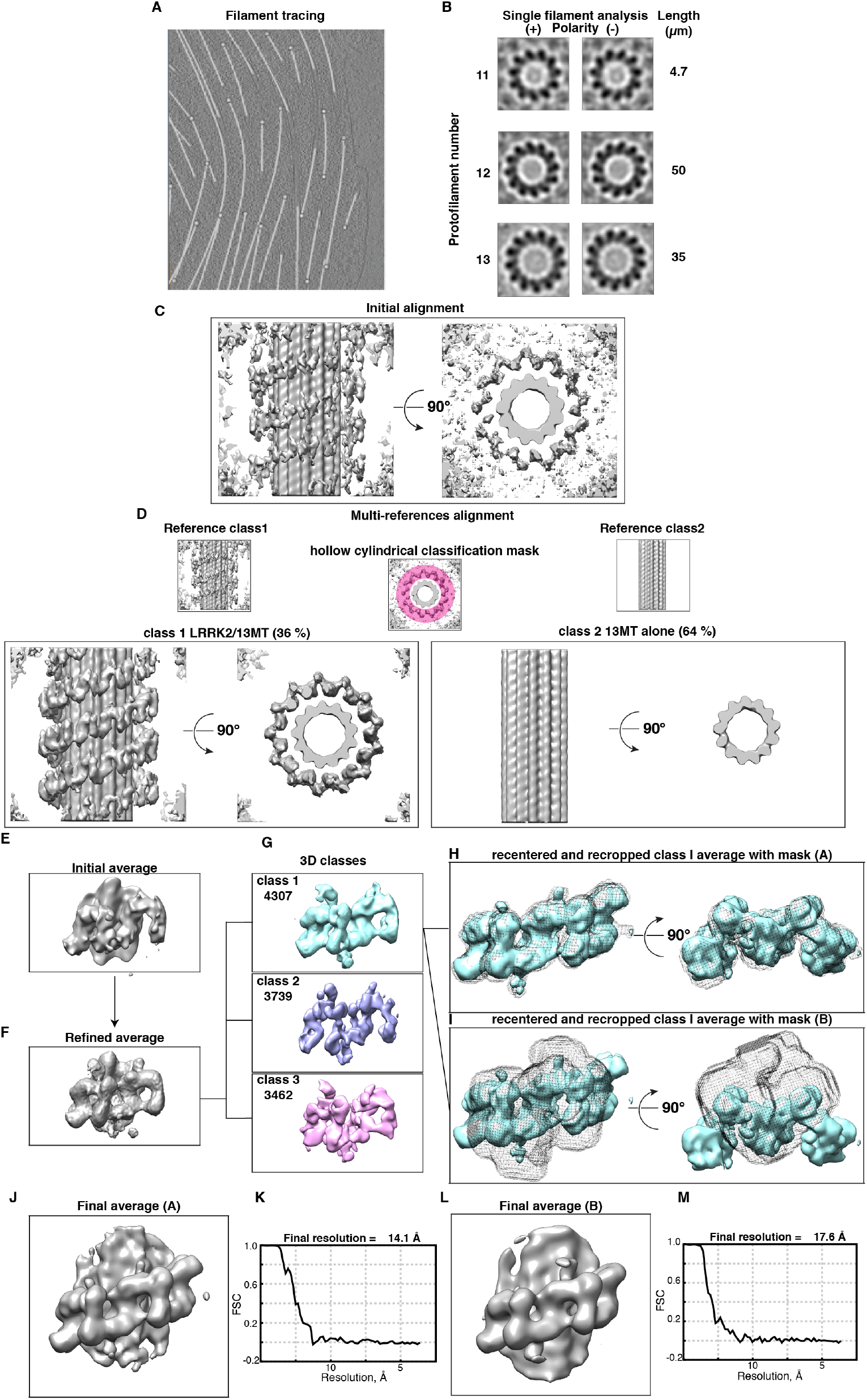
Subtomogram averaging. (A) Individual microtubules are automatically annotated using the filament tracing function of Amira (TFS). The annotated MTs (grey) are indicated in a slice of a tomogram showing LRRK2-decorated MT. (B) Single microtubule averaging sampled at every 4 nm was performed using Dynamo to determine the protofilament number and polarity of individual MTs using 4 binned tomograms. Examples of a cross-section of averages from the plus end (anti-clockwise slew, left column) and from the minus end (clockwise slew, right column). The total length of each class of MTs unambiguously determined from LRRK2 tomograms is shown. (C) After sorting of MTs according to the protofilament number, larger subtomograms (70 nm) were re-extracted and averaged to obtain initial LRRK2/MT averages. (D) Dynamo multi-reference alignment using two different templates: (1) LRRK2/MT and (2) MT only, using a hollow cylindrical classification mask corresponding to LRRK2 density, resulting in averages for LRRK2/13-protofilament MT (class 1) and non-decorated 13-protofilament MT (class 2). The same protocol was followed for 11-pf and 12-pf MTs. (E) Helical parameters of the LRRK2 helix on 12-protofilament MT were estimated and used for extraction of subtomograms containing the LRRK2 protomer from un-binned and non CTF-corrected tomograms and used to create an initial average using pre-assigned orientations. (F) 3D refinement of extracted particles was performed in RELION to create a template for subsequent classification. (G) The refined average from (F) was used as a reference for 3D classification into three classes in RELION. Classes with high levels of noise (classes 2 and 3) were excluded. (H, I) Particles belonging to the good class (class 1) were subjected to gold-standard refinement and post-processing in RELION using two different masks shown in grey (A and B). (J, L) The final subtomogram averages from LRRK2/12-protofilament MT. (K,M) FSC curves for the two maps shown in J and L, respectively.

**Fig. S2.**
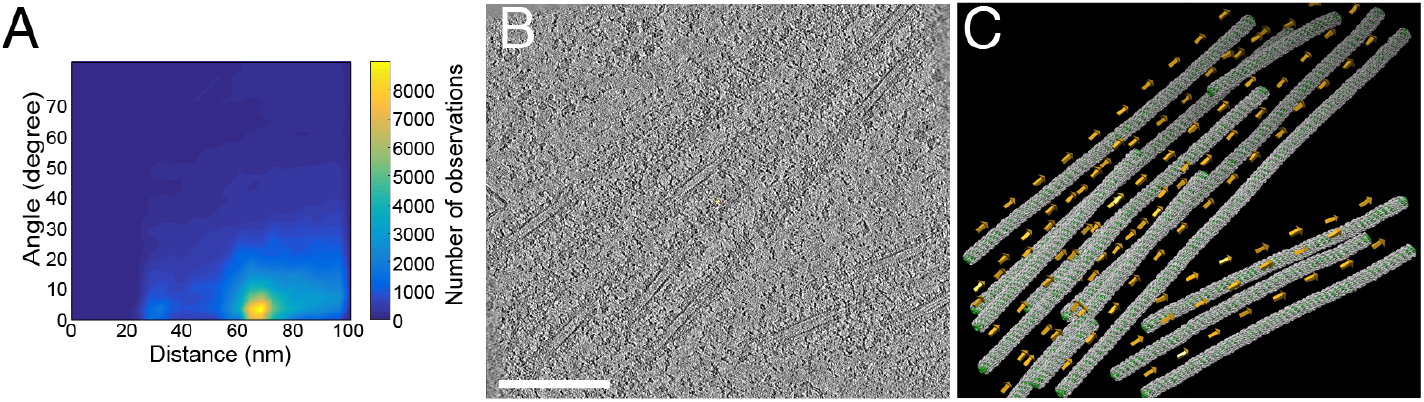
LRRK2-decorated MTs are found in bundles with consistent polarity. (A) Relative angle and distance between MT segments revealing parallel bundles with a regular center-to-center distance of 70 ±18 nm (peak ∼67 nm) between microtubules. (B) Slice of a tomogram showing bundles of LRRK2-decorated MT. Sale bar, 200 nm. (C) Individual MT analysis with Dynamo shows that MTs found in bundles have consistent MT polarity.

**Fig. S3.**
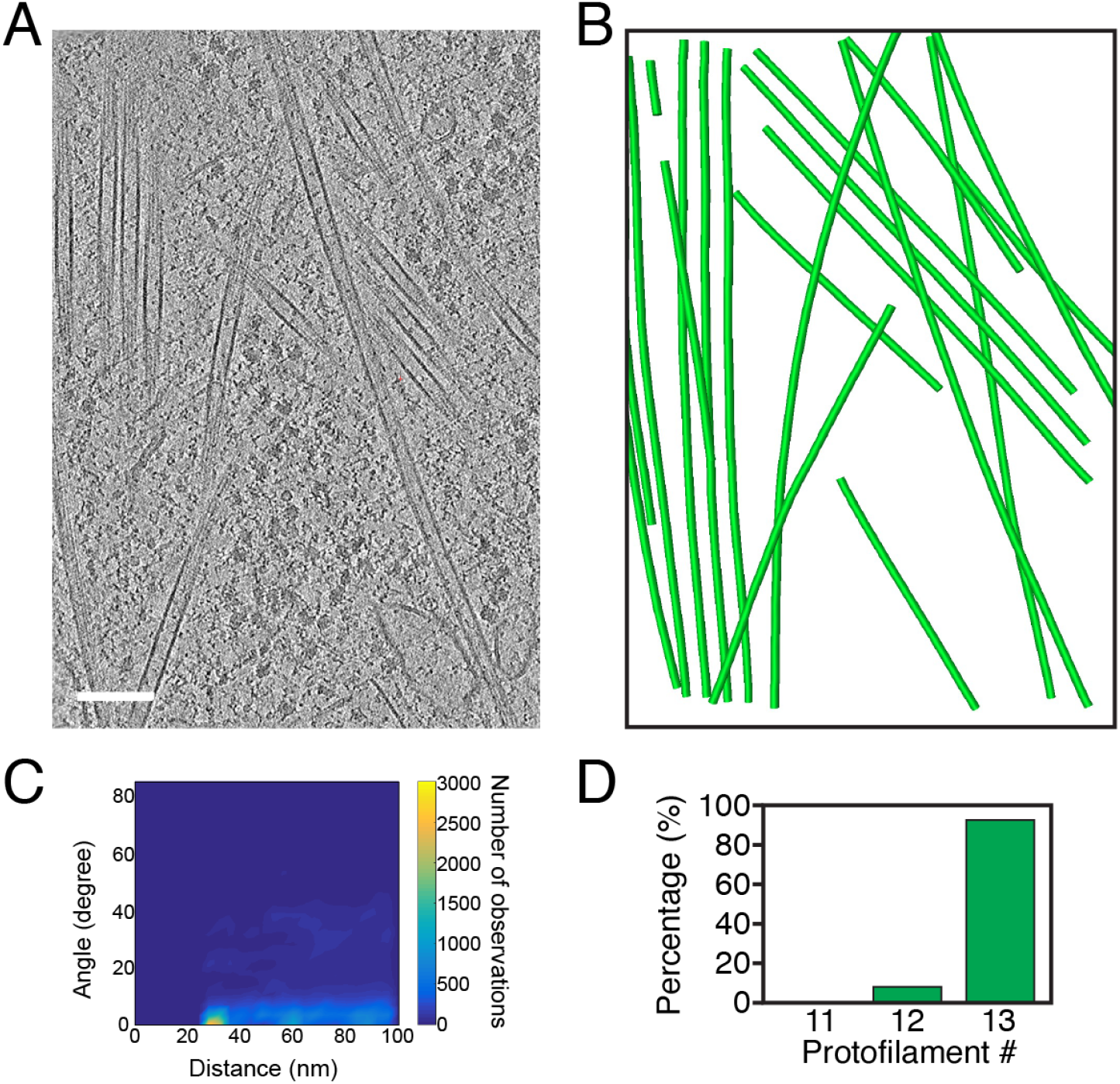
MTs in Taxol-treated HEK-293T cells. (A) A slice of a tomogram showing MTs in HEK-293T cells treated with 5 *μ*M Taxol. Scale bars, 100 nm. (B) The annotated MTs (in green) in the corresponding tomogram. (C) Relative angle and center-to-center distance between MT from 5 Taxol-treated cellular tomograms. The average distance is 47 ± 32 nm (peak ∼31 nm). (D) Graph showing the percentage of different MT classes found in 5 tomograms of Taxol-treated cells from two biological duplicates.

**Fig. S4.**
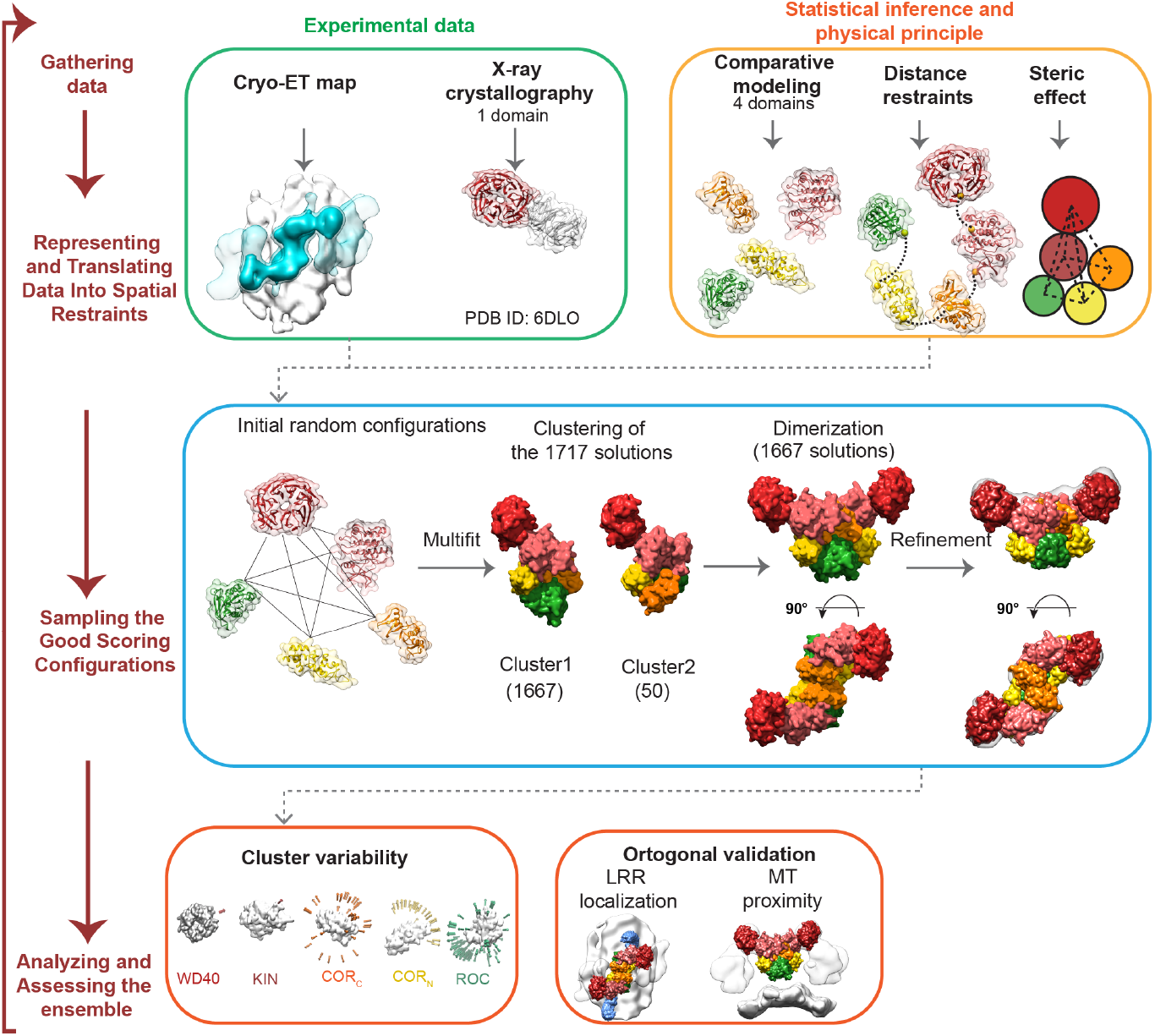
Integrative structure determination of the domain architecture of LRRK2. First, we gather data generated by various experiments and computational methods. Second, the five C-terminal domains of LRRK2 are represented as rigid bodies and the data are translated into spatial restraints. Third, an ensemble of structures that satisfy the restraints is sampled using Multifit, an IMP module that allows simultaneously fitting the rigid bodies respecting the a priori restraints imposed. The ensemble is clustered into distinct domain arrangements and representative structures are further refined. Finally, the refined models are analyzed for accuracy and certainty (See Materials and Methods for details).

**Fig. S5.**
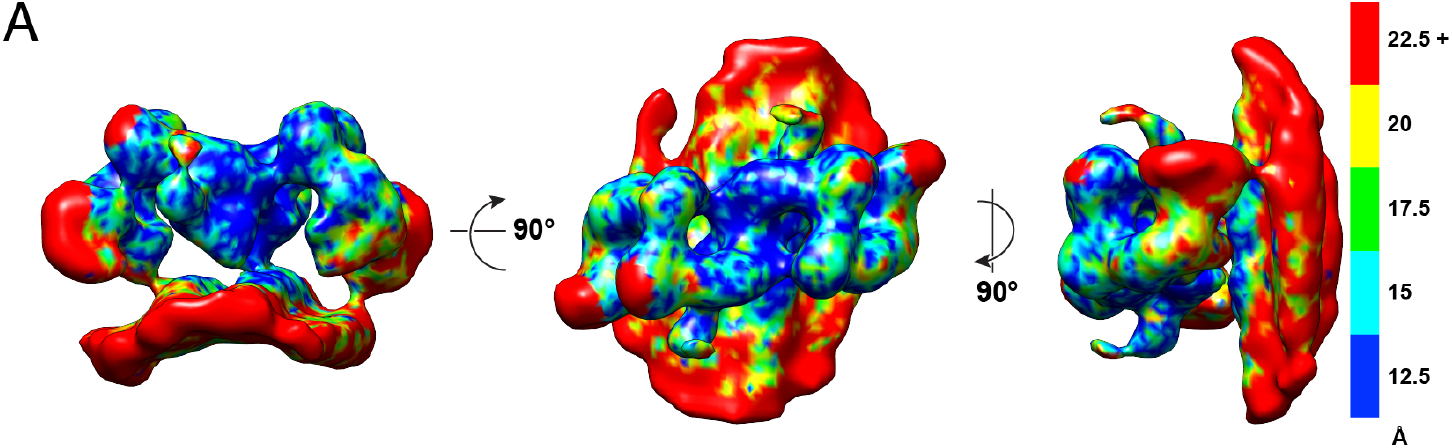
Local resolution of subtomogram-average 3-D reconstructions. Local resolution for the map shown in Fig. 3G.

**Fig. S6.**
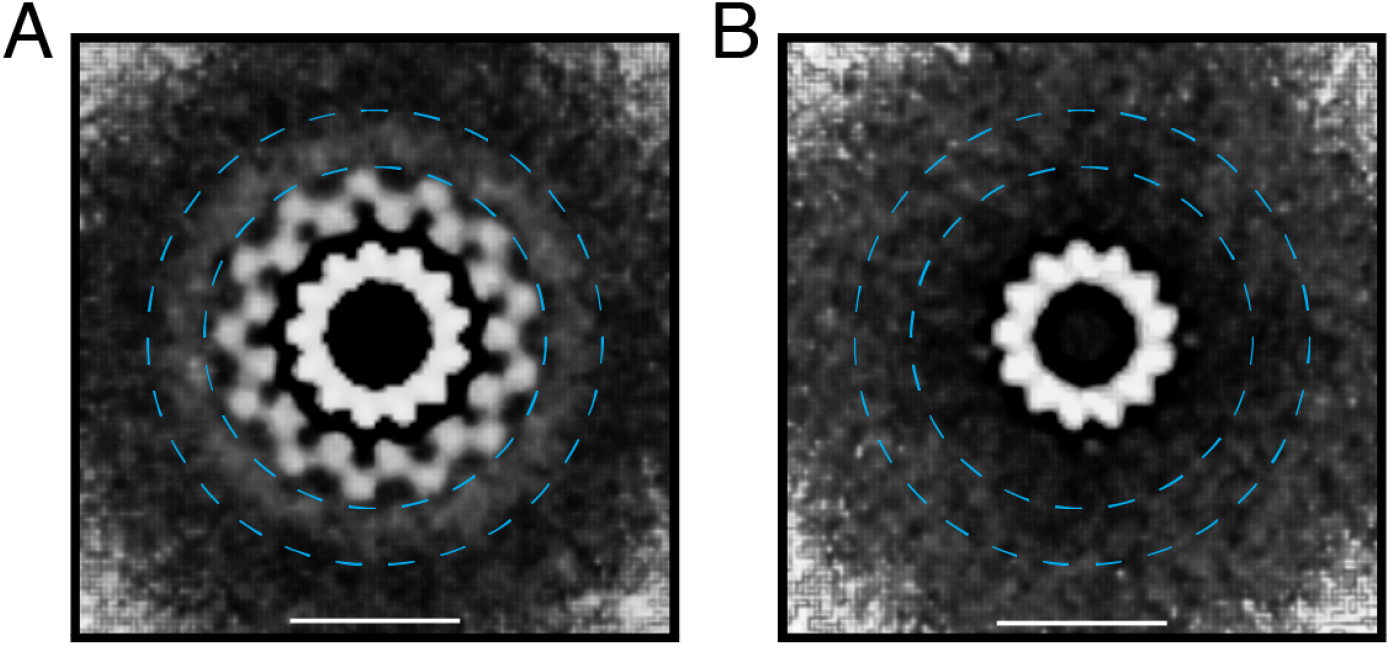
Putative location of LRRK2’s N-terminal domains. (A) The cross-section density view of LRRK2/13-pf MT at lower threshold. The weak outermost density highlighted with a pair of blue dotted circles is observed. (B) The cross-section density view of an undecorated 13-pf MT at lower threshold. No outer density is observed in this cross-section. Two pairs of dotted blue circles shown in A and B correspond to 50 nm and 67 nm in diameter, respectively. Cf. the peak of center to center distance between LRRK2-decorated MT bundles is ∼67 nm (Fig. S2A). Scale bar: 25 nm.

**Fig. S7.**
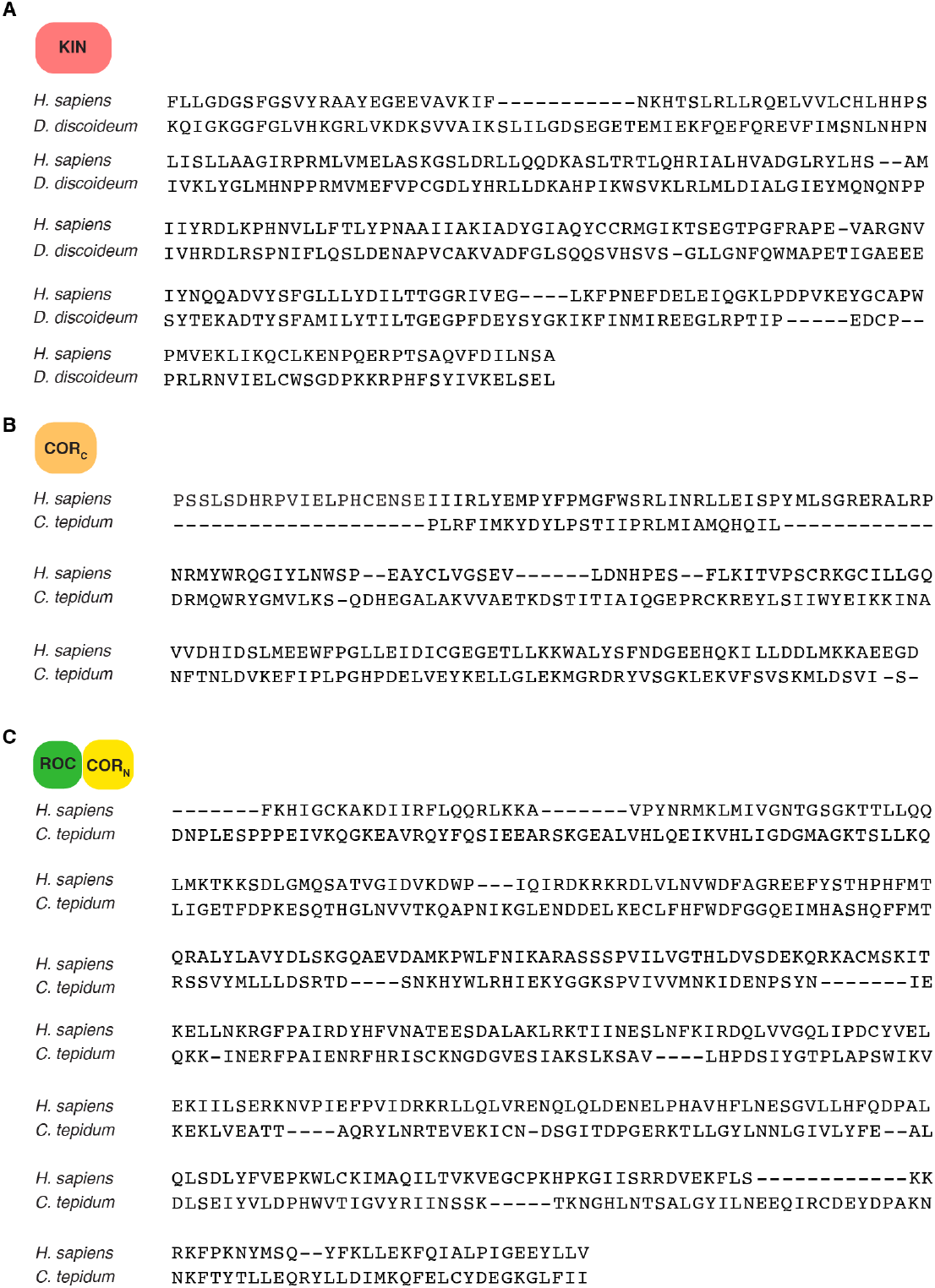
Sequence alignment used for homology modeling of LRRK2. (A) Alignment of the LRRK2 kinase domain to the *D. discoideum* Roco4 kinase domain template structure PDB ID code 4F0F (bound to AppCp). (B) Alignment of the LRRK2 COR_C_ (C-terminal end of ROCCOR) domain to the *C. tepidum* Roco ROCCOR template structure PDB ID 3DPU. (C) Alignment of the LRRK2 COR_N_ and ROC to the *C.tepidum* Roco ROCCOR template structure PDB ID 3DPU.

**Fig. S8.**
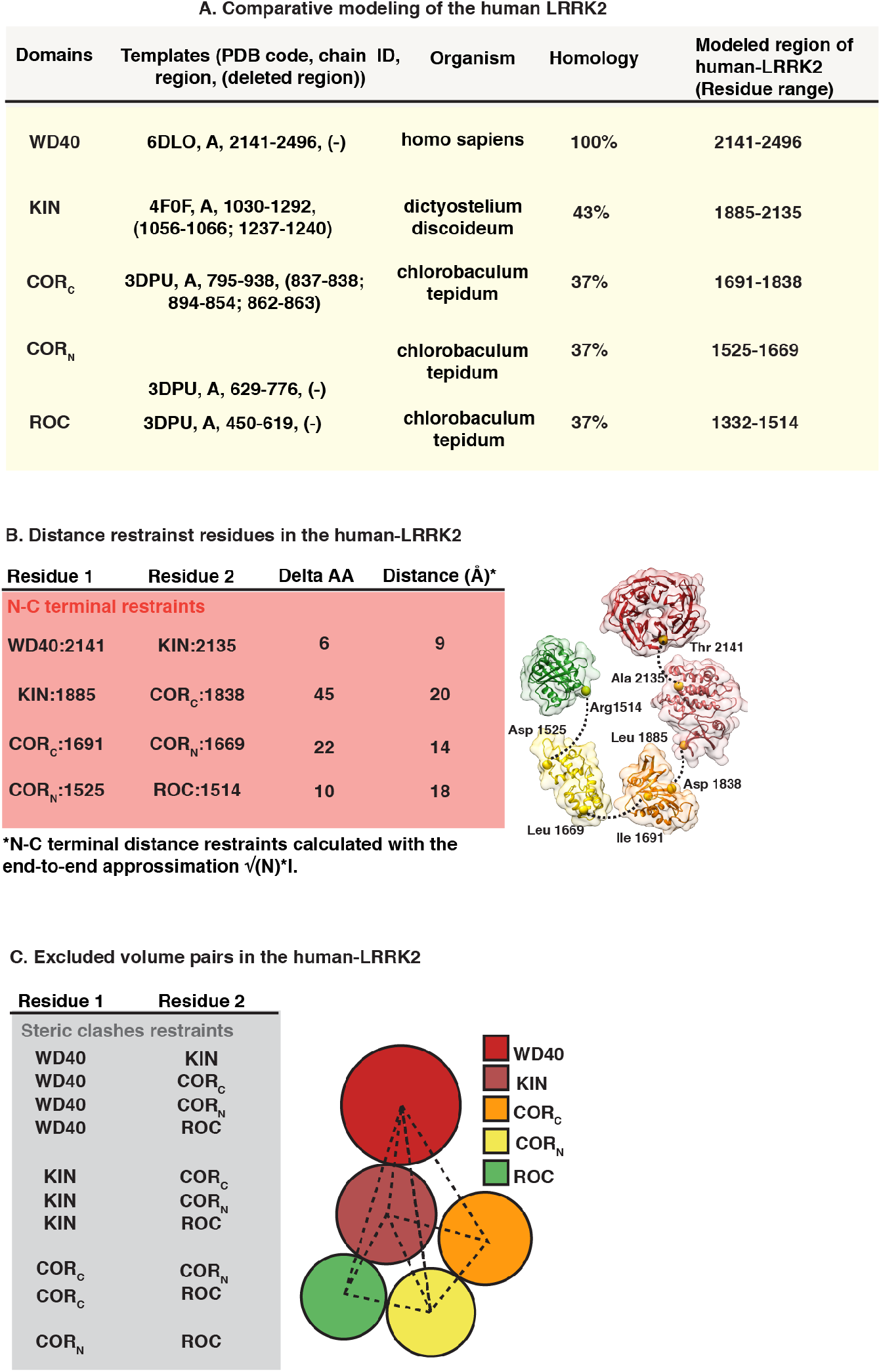
Experimental data, statistical inference and physical principles considered in the integrative modeling procedure. (A) Comparative modeling templates for WD40, KIN, COR_C_, COR_N_, and ROC. The PDB template, the organism, the homology, and the relative modeled region are reported for each domain. (B) Distance restraints and (C) exclude volumes pairs considered in the integrative modeling approach.

**Fig. S9.**
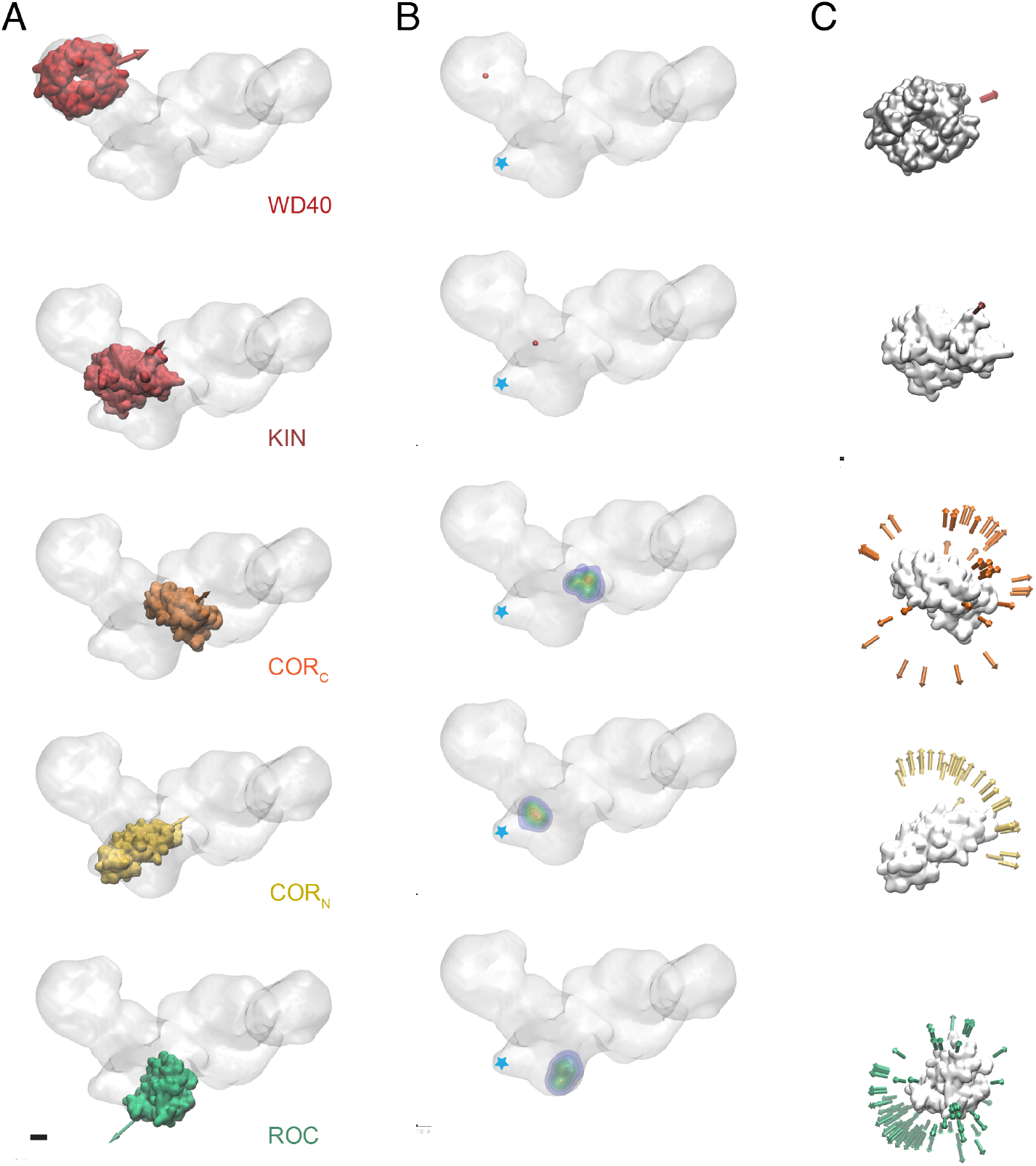
Ensemble Variability Analysis. Analysis of the ensemble variability for WD40 (red), KIN (pink), CORC (orange), CORN (yellow) and ROC (green). (A) Representative structure from the ensemble, with the position and orientation indicated by an arrow. (B) Probability density function of the localization of the centroid of each domain within the density. A blue asterisk indicates the potential location of the C-terminus of the LRR domain. (C) Orientation variability analysis of each domain within the ensemble; each arrow indicates the orientation of the domain with respect to their center of mass colored accordingly to the domain analyzed. Scale bar: 5 nm.

**Table S1.**
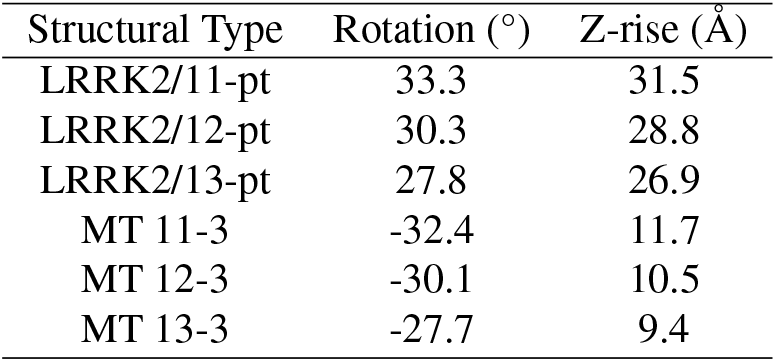
Parameters for the LRRK2 helices and MTs found in cells.

**Table S2.**
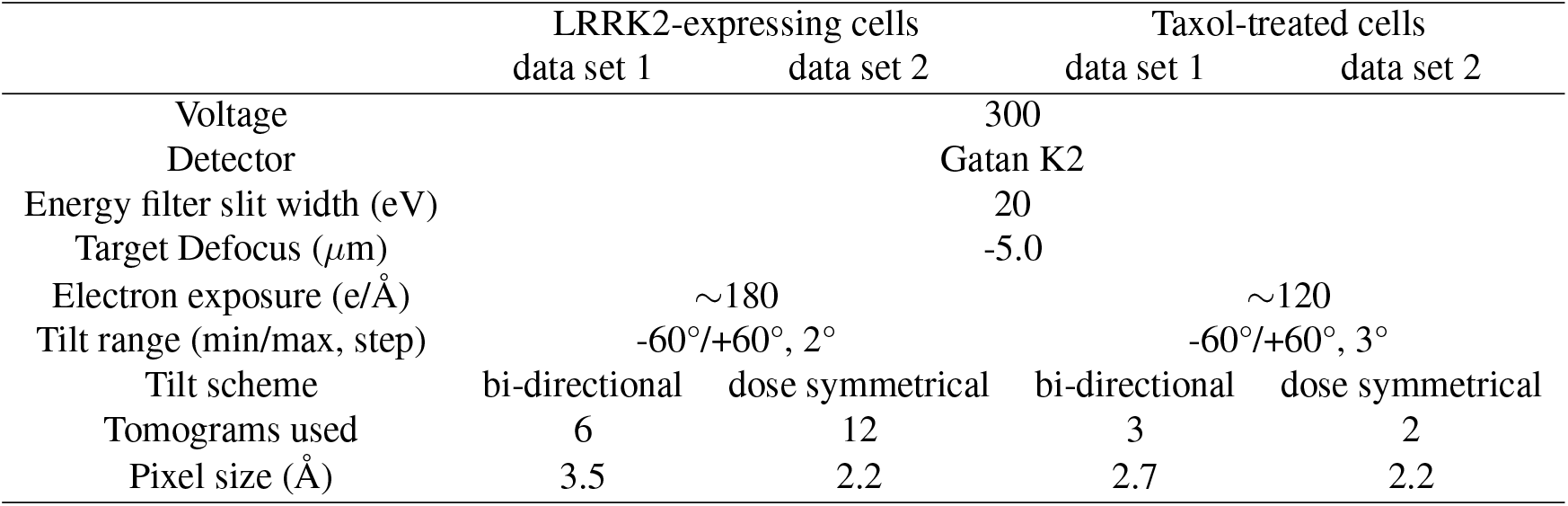
Cryo-ET data collection.

